# Mitochondrial Deletions and Healthspan: Lessons from the Common Repeat

**DOI:** 10.1101/2024.09.02.610808

**Authors:** Victor Shamanskiy, Konstantin Gunbin, Evgenii O. Tretiakov, Ilya Mazunin, Viktoria Skripskaya, Alina A. Mikhailova, Alina G. Mikhailova, Natalia Ree, Valeriia Timonina, Dmitry Knorre, Wolfram S. Kunz, Nathan Fiorellino, Alexandre Reymond, Georgii A. Bazykin, Jacques Fellay, Masashi Tanaka, Konstantin Khrapko, Konstantin Popadin

## Abstract

Aging, characterized by a series of functional declines correlated with advancing chronological age, has a significant mitochondrial DNA (mtDNA) component, with somatic mtDNA deletions playing a central role. In post-mitotic or slow-dividing cells like neurons and skeletal muscles, selfish mtDNA deletions clonally expand within a cell, ultimately leading to the deterioration and death of host cells and appearence of age-related phenotypes. Thus reducing the burden of somatic deletions could have far-reaching systemic benefits for the entire human body. Given the crucial role of direct nucleotide repeats in the formation of mitochondrial deletions, we hypothesize that minimizing these repeats in the human mitochondrial genome could enhance healthspan by decreasing somatic deletions. To investigate this hypothesis, we focus on the “common repeat”, a 13-base pair perfect direct repeat sequence (ACCTCCCTCACCA) located at positions 8470-8482 and 13447-13459, respectively. This perfect repeat: (i) is highly prevalent, with its potential deleterious consequences affecting the majority of humans; (ii) represents one of the most fragile sites, highly prone to forming deletions; (iii) when disrupted, is associated with a decreased somatic deletion load and enhanced human healthspan; (iv) is likely to experience positive selection in the present or near future due to indirect fitness effects, such as the “grandmother effect”, and direct fitness effects, such as (v) a decreased mutation rate. These observations support the argument that reducing the mtDNA somatic deletion load through targeted disruption of these repeats, or by using naturally occurring polymorphisms with disrupted repeats in mitochondrial medicine, could be an effective approach to increasing human longevity.

## Introduction

Aging encompasses a series of functional declines that correlate with advancing chronological age, typically beginning after sexual maturity [(Melzer et al. 2020)]. Genetically, aging can be explained by the inability of natural selection to eliminate harmful traits that manifest themselves after reproduction [(Fisher, 1930), (Medawar, 1946), (Medawar, 1952), (Williams, 1957), (Hamilton, 1966)]. These traits, known as Deleterious In Late Life (DILL) alleles [(Cortopassi, 2002)], persist because natural selection mainly focuses on preserving fitness during the reproductive phase and is less effective in the later stages of life. Some DILL alleles may also have Beneficial In Early Life (BIEL) effects, which can increase their presence in the population. While the complexity of aging has led to the use of whole-genome approaches [(Melzer et al., 2020)], studying individual DILL alleles is crucial for gaining specific insights into the mechanisms of aging. This focus also helps clarify the limitations of population-based whole-genome studies for investigating aging and other complex traits (see discussion).

DILL alleles are germ-line variants, which can cause increased somatic genome instability with age. Such DILL alleles can originate either in the nuclear or mitochondrial genome. Mitochondrially-encoded DILL variants, despite the small size of mtDNA, are expected to play a significant role in aging due to the smaller effective population size of mtDNA versus nuclear DNA, which increases the likelihood of fixation of all types of slightly deleterious variants, including DILL alleles. After the origin of the mitochondrially-encoded germline DILL allele it can be very effectively realized in mtDNA - i.e. lead to deleterious somatic mutations with age-related consequences - due to (i) high mtDNA somatic mutation rate, during which a somatic mutation can originate and (ii) semi-autonomous nature of mtDNA replication, allowing the somatic mtDNA mutation to propagate within a cell, leading to age-related phenotypes. Among the various somatic mutations in mtDNA, deletions are strongly linked to aging and are commonly found in aged post-mitotic tissues [(Cortopassi, 2002), (Kraytsberg et al., 2006), (Lujan et al., 2020), (Trifunovic et al., 2004)]. Origin and clonal expansion of the majority of somatic mtDNA deletions till the phenotypically important threshold takes dosens of years, eventually leading to host cell degeneration and age-related conditions in the late life. Thus, somatic mtDNA deletions work as effective and important immediate contributors to ageing process, and understanding DILL alleles behind of them - alleles which predispose the origin of these somatic mtDNA deletions - is of great importance. Deletions can occur due to defects in mtDNA replication machinery [(Trifunovic et al., 2004), (Lujan et al., 2020)] or due to the inherent fragility of the mtDNA structure [(Shamanskiy et al., 2023); (Guo et al., 2010)]. It is highly possible that contribution of both - defective mtDNA replication machinery and highly fragile structure of mtDNA can interact synergistically leading to strongly increased aging [(Hinton et al., 2024)], however this type of crosstalk between nuclear and mitochondrial DILL alleles are beyond the scope of the current paper and in this study we focus on the variation in the abundance of the direct repeats in human mtDNA, which are considered as DILL alleles.

Current in vivo, in vitro [(Persson et al., 2019)], and in silico [(Shamanskiy et al., 2023)] models indicate that direct nucleotide repeats are crucial in the formation of mitochondrial deletions. Most human mtDNA deletions are indeed flanked by direct repeats or repeat-like sequences [(Guo et al., 2010)]. Additionally, comparative species studies reveal a negative correlation between the abundance of direct nucleotide repeats in mtDNA and lifespan in mammals [(Khaidakov et al. 2006), (Samuels, 2004)]. This suggests that a high burden of mitochondrial deletions associated with these repeats may limit lifespan. Based on these insights, we hypothesize that reducing the number of direct repeats within the human mitochondrial genome could enhance human healthspan by decreasing the burden of somatic mtDNA deletions. To investigate this hypothesis, we focus on the “common repeat,” a 13-base pair perfect direct repeat sequence (ACCTCCCTCACCA) located at positions 8470-8482 and 13447-13459 (Figure 1A). We demonstrated that this perfect repeat: (i) is highly prevalent, thus its potential deleterious consequences affect the majority of humans; (ii) represents one of the most fragile sites, highly prone to forming deletions; (iii) when disrupted, is associated with a decreased somatic deletion load and enhanced human healthspan; (iv) when disrupted is likely to experience positive selection in the present or near future due to indirect fitness effects, such as the “grandmother effect,” and direct fitness effects, such as (v) a decreased mutational rate.

**Figure 1.**
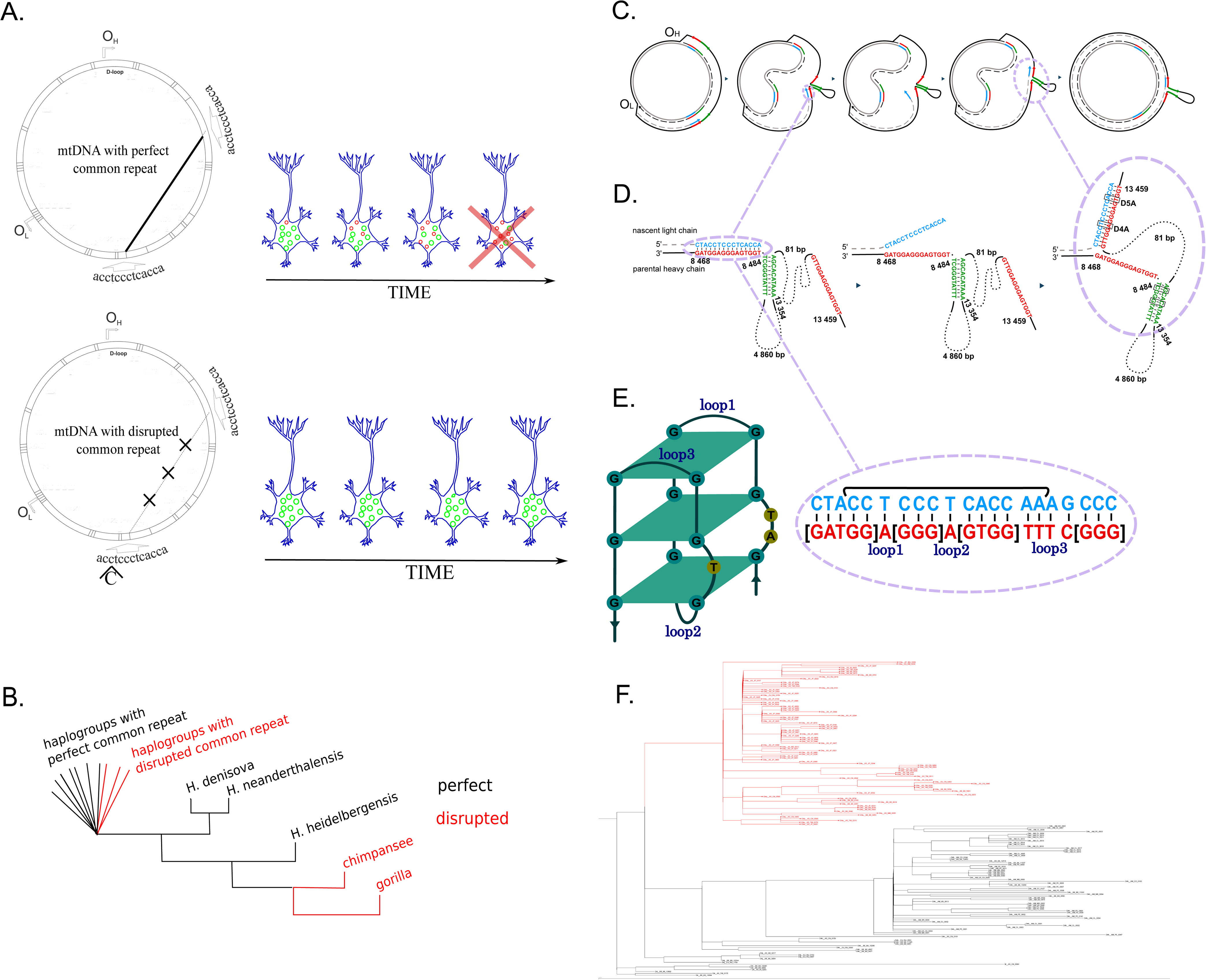
**a.** Working hypothesis: disrupted common repeat impedes the origin of somatic mtDNA deletions, maintaining postmitotic cells (neurons and skeletal muscular cells) in more healthy conditions (green circles – wild type mtDNA; red small circles – mtDNA with common deletion), thus postponing age-related phenotypes. **b.** The phylogenetic scheme of the common repeat. Perfect common repeat is human-specific **c.** The replication slippage can explain mtDNA common deletion formation, where nested inverted (green) and direct (red) repeats play a crucial role during the replication of the lagging (parental heavy chain) strand. Main steps of the deletion formation. First, according to the asymmetric replication of mtDNA, the lagging strand (solid black line) spent a significant amount of time in the single-stranded condition. Second, a single-stranded parental heavy chain forms a secondary structure shaped by inverted repeats. This stem can arrest the replication fork. Third, replication arrest leads to partial dissociation of the newly synthesised proximal arm of the direct repeat (blue arrow). Fourth, the dissociated chain realigns to the closest distal arm of the direct repeat. Fifth, replication has resembled, but the newly-synthesized light chain (dotted grey line) has a deletion - both inverted repeats and DNA between them as well as the proximal arm of the direct repeat are removed; **d.** Common deletion formation on nucleotide scale. First: synthesis of the nascent light chain (blue) and arrest of the replication near the stem, formed by the inverted repeat. Second: partial dissociation of the nascent proximal arm of the common repeat. Third: realigning of the newly-synthesized proximal arm of the common repeat with the distal arm of the common repeat and continuation of the replication. The most frequent polymorphisms (with 100 or more mtDNAs from HmtDB) disrupting these repeats are marked by rectangles. These substitutions can substantially slow down the process of deletion formation. The common repeat here is extended from a classical perfect 13bp-long repeat (ACCTCCCTCACCA) to a 15bp-long degraded repeat with one mismatch (CtACCTCCCTCACCA). **e.** Scheme of the GQ, overlapped with the first arm of the common repeat **f.** The simplified phylogenetic tree of D4a haplogroup (red) with neighbour branches (black). The shortened D4a-specific branches can be explained by decreased mtDNA mutation rate.

## Results

### 1. Mitochondrial common repeat is perfect in more than 98% of the human population

Analysis of complete mitochondrial genomes from the HmtDB database ([(Clima et al., 2017)]; N = 43,437) reveals that the common repeat is intact (i.e. is perfect) in more than 98% of all individuals (42,641 out of 43,437). Disruptions have occurred independently several times in relatively recent haplogroups (Supplementary Table 1; Figure 1B). The most frequent disruption, m.8473T>C, is found in haplogroups including D4a, R2, U2e, H1c, and U6a, each supported by over 20 genomes with this variant. The second most common disruption, m.8472C>T, is prevalent in the Ashkenazi-specific haplogroup N1b, and the third, m.8479A>G, occurs in the D5a haplogroup. Further analysis using the HelixMTdb database ([(Bolze et al., 2020)]; N = 196,554) confirms the rarity of disrupted common repeats: m.8473T>C is found in 1% (2,118 out of 196,554) and m.8472C>T in 0.4% (833 out of 196,554) of genomes. Therefore, since the common repeat remains intact in the vast majority of human genomes, any deleterious effects associated with this sequence (see chapter 2) could potentially impact nearly the entire human population.

### 2. Mitochondrial common repeat is one of the most fragile repeat in the human mtDNA

The common repeat exhibits particular fragility, defined as a propensity to form deletions, for several reasons (Figures 1C-1E): (i) it is associated with the most frequent and pathogenic deletions, both somatic and germline, identified in human mtDNA; (ii) it represents the longest perfect direct repeat in the major arc of human mtDNA; (iii) it is located within a “contact zone” where the probability of interaction between the two arms of the repeat is maximized due to their spatial proximity; (iv) nested within the arms of the common repeat there is an inverted repeat, which, if folded, can pause replication and effectively reduce the distance between the arms; (v) the first arm of the common repeat is embedded within a robust G-quadruplex, which can further stall the replication fork, heightening the risk of deletions (Figure 1E); (vi) the sequence of the common repeat closely resembles a well-known hotspot sequence that facilitates recombination. Below we discuss all these properties of the common repeat in more details.

i. Empirically, the common deletion - deletion flanked by the arms of the common repeat - has been linked to a spectrum of rare diseases, such as progressive external ophthalmoplegia [(Moraes et al., 1989)], Kearns-Sayre syndrome [(Zeviani et al., 1988), (Moraes et al., 1989)], as well as more common complex phenotypes including parkinsonism [(Bender et al., 2006)] and other aging-related conditions [(Kraytsberg et al., 2006)]. It is identified in numerous studies as one of the most frequent and abundant mtDNA rearrangements [(Samuels et al. 2004)], marking it as a vital indicator of mitochondrial deletion load within patients and across various cells or tissues. Recent advancements in deep NGS analysis of DNA [(Lujan et al., 2020)] and RNA [(Omidsalar et al., 2024)] demonstrate that although this deletion may not always be the most prevalent across different human tissues, it still represents a significant portion that profoundly influences human phenotypes.
ii. The mutagenic properties of direct repeats are known to increase exponentially with their length. This correlation was first established through bacteriophage deletion experiments [(Pierce et al. 1991)] and was subsequently validated on a comparative-species level by Khaidakov et al. They calculated the total mutagenicity scores of mtDNA for each species by evaluating each individual direct repeat in the mtDNA, factoring in the effects of length, and found a strong negative correlation between these aggregated scores and mammalian lifespan [(Khaidakov et al., 2006)]. Given that the 13-base-pair common repeat is the longest perfect direct repeat in the major arc of human mtDNA, it is expected to exhibit the highest mutagenic potential.
iii. Through analysis of human mtDNA structure during replication, we developed a model in which the single-stranded parental heavy chain of the major arc forms a substantial hairpin-like loop, featuring contact zones between 6–9 kb and 13–16 kb [(Shamanskiy et al., 2023)]. Direct repeats within this contact zone, due to their spatial proximity, show a threefold increase in deletion activity. Specifically, the common repeat, positioned with its first arm at 8470–8482 bp and its second arm at 13,447–13,459 bp, lies within this contact zone, significantly enhancing its susceptibility to deletions.
iv. In addition to the global contact zone, the local secondary structure of the single-stranded parental heavy DNA can be stabilized by microhomologies such as inverted repeats, which may hybridize to form stem structures [(Shamanskiy et al., 2023)]. Our analysis of the region adjacent to the common repeat has revealed nested inverted repeat (see green colored sequences in Figure 1D) that could both pause the replication fork and reduce the effective distance between the arms of the common repeat. This shortened distance enhances the likelihood of deletions by promoting closer interaction between the arms of the common repeat (Figures 1C-1D).
v. Numerous studies have linked G-quadruplexes (GQs) to mtDNA deletions [(Dong et al., 2014), (Doimo et al., 2023)]. The heavy strand of mtDNA, which is enriched with GQs, may experience pauses in polymerase activity when encountering these structured regions. This interruption increases the likelihood of dissociation of the newly synthesized daughter strand initiating the deletion formation (see Figures 1C-1D for the proposed mechanism of common deletion formation). Our research examined whether the first or the second arm of the common repeat is associated with a GQ. Using data from both in vitro [(Bedrat et al. 2016)] and in silico studies of GQs [(Hon et al. 2017)], we observed that the first arm of the repeat (8470– 8482 bp) is overlapped with a stable GQ (see Figure 1E). It is important to highlight that completely in line with our findings, according to the current “copy-choice recombination” model of deletion formation [(Persson et al., 2019), (Shamanskiy et al., 2023)] specifically the first—not the second—arm of the direct repeat should be responsible for pausing replication (see also Figures 1C,1D).
vi. A notable characteristic of the common repeat is its nucleotide sequence (aCCTCCCTCACCAx), which closely mirrors the degenerate 13 bp motif (CCNCCNTNNCCNC) commonly found at human nuclear recombination hotspots and aligns with the predicted binding domain for the PRDM9 protein [(Berg et al., 2021), (Muñoz-Fuentes et al. 2011), (Tanaka et al. 1998)]. After years of debate and uncertainty regarding rare mtDNA recombination events in humans and mammals, these phenomena have now been definitively confirmed through various experimental approaches [(Fragkoulis et al., 2024)]. These findings suggest that such recombination events generally adhere to the “copy-choice recombination” model, which is also implicated in the mechanism of deletion formation. This research shows that mammalian mtDNA recombination often involves replisome-mediated template switching among microhomologous sequences, either between different molecules, which can initiate recombination, or within the same molecule, leading to deletions (Figures 1C-1E). Although intermolecular recombination is exceedingly rare in stable tissue conditions—requiring interaction between two mtDNA molecules with different sequences and thus necessitating heteroplasmic nucleoids within mitochondria—intramolecular recombination among microhomologous sequences is a common source of pathological mtDNA deletions and other rearrangements. The recurring presence of the CCNCCNTNNCCNC motif at hotspots for both recombination and deletion suggests a pivotal, though not fully understood, role in these genetic dynamics.

Altogether, the six traits of the common repeat discussed above establish it as one of the most fragile regions in human mtDNA, playing a crucial role in genomic instability and susceptibility to deletions.

### 3. Mitochondrial common repeat loses fragility when disrupted: decreasing somatic deletions and enhancing human longevity

The factors outlined in Chapter 2 illustrate the high fragility of the perfect common repeat to form deletions. By altering this repeat through rare nucleotide substitutions (see Chapter 1), its structure becomes imperfect or degraded, potentially reducing its fragility and lowering the likelihood of both germline and somatic deletions. Decreased incidence of germline deletions is expected to lead to decreased incidence of rare diseases such as progressive external ophthalmoplegia [(Moraes et al., 1989)], Kearns-Sayre syndrome [(Zeviani et al., 1988), (Moraes et al., 1989)] and Pearson syndrome [(Yoshimi et al. 2022)]. Due to the rarity of such conditions, haplogroup-specific analysis requires more data to test this hypothesis, especially from non-European populations (for example, samples rich in D4a from Eastern populations). In turn, some consequences of the decreased incidence of somatic deletions in samples with disrupted common repeat is easier to test using available data. For example, we expect that disrupted repeat would correlate with (i) a decreased somatic burden of deletions, (ii) decreased incidence of complex age-related diseases with a mitochondrial component such as neurodegeneration and sarcopenia which altogether contribute to increased lifespan.

To test the decreased incidence of the somatic mtDNA deletions in samples with disrupted common repeat we analyzed frontal cortex samples from two elderly individuals [(Guo et al., 2010)] in the Ashkenazi-specific haplogroup N1b [(Costa et al., 2013), (Feder et al., 2008)], which carries the germline variant m.8472C>T. Originally, when comparing the global deletion spectra between two samples with disrupted common repeats and two control samples, we did not observe any significant overall shifts [(Guo et al., 2010)] (see also Fig 2 left panel). This indicated that the global pattern of deletions remained similar across the groups [(Guo et al., 2010)], characterized, for example, by two peaks of breakpoints: one around position 8000 (5’) and another around position 13000 (3’) on the mtDNA. At the time, we interpreted this finding as contradictory to the hypothesis that the common repeat is responsible for the majority of mtDNA deletions [(Samuels et al. 2004)], i.e., it shapes the global pattern of deletions. Currently, we are focusing on the local effects of the disrupted common repeat. We hypothesize that such disruptions would not alter the global deletion pattern but would lead to a significant decrease in the number of common deletions specifically. To test this, we reanalyzed the data from [(Guo et al., 2010)] (see Methods) and found that, indeed, in the local region of the common repeat, there is an approximately tenfold deficit in deletions in cases (with disrupted repeats) compared to controls (see Figure 2 right panel). This finding supports the notion that disrupting the 13 bp repeat even by one mismatch reduces the formation of common deletions drastically. Interestingly, authors of a recent study, which demonstrated that mtDNA deletion formation results from copy-choice recombination [(Persson et al., 2019)], also found that disruption of any arm of the common repeat prevents deletion formation in vitro [(Persson et al., 2019)]. Similarly, bacterial deletion frequency declines tenfold when direct repeats contain single nucleotide mismatches [(Albertini et al. 1982)]. Altogether, these findings corroborate our hypothesis that disruption of the common repeat significantly reduces the burden of common deletions.

**Figure 2.**
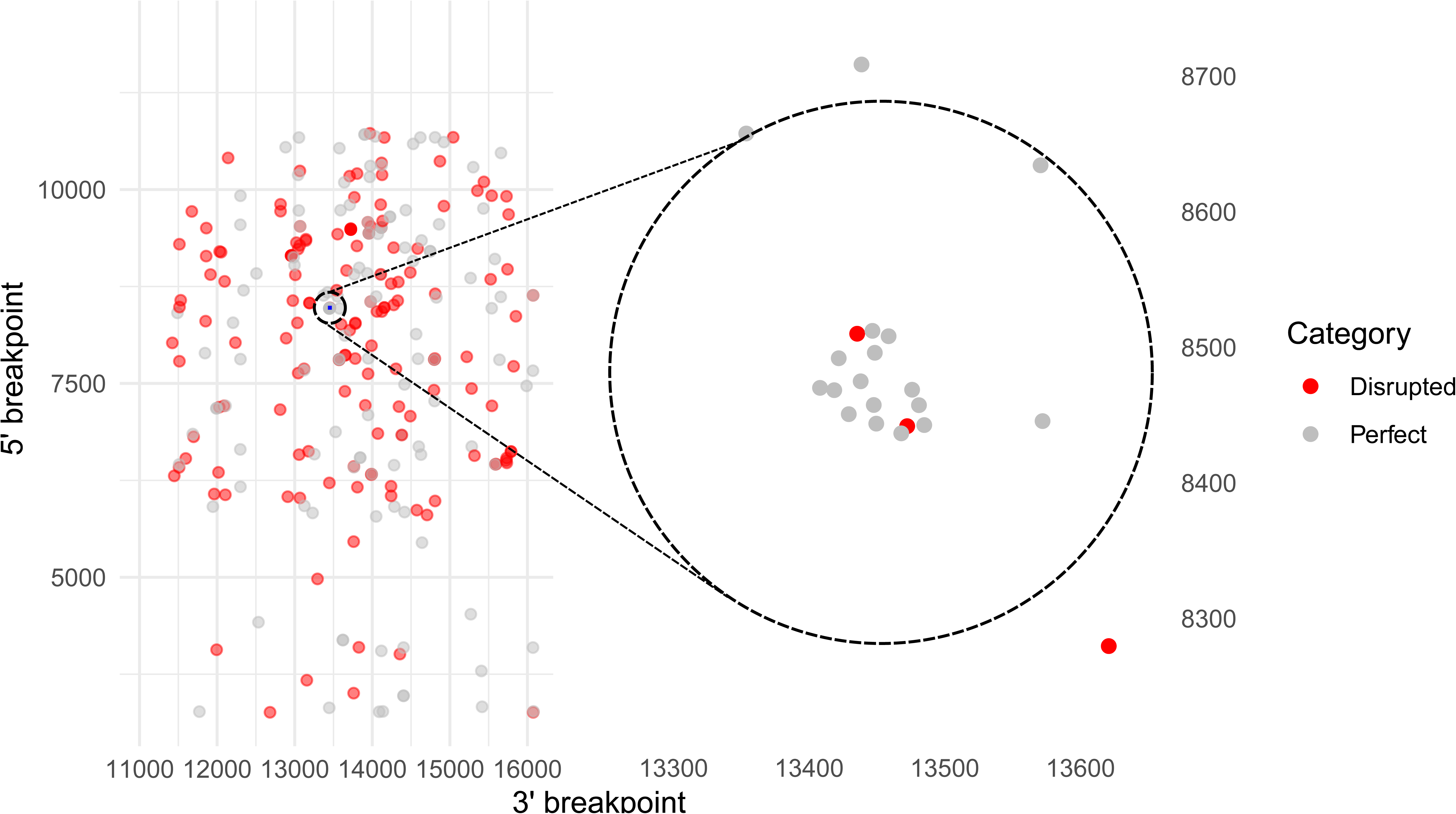
Distribution of deletion breakpoints in frontal cortex samples with a perfect common repeat (grey) and disrupted common repeat (red). While the global deletion patterns (left panel) appear similar, a local analysis within a defined region around the common repeat (centered at 13,453 and 8,476 bp, radius = 200 bp, indicated by the circle) reveals a significant reduction in deletions in samples with a disrupted common repeat (right panel: 2 vs. 17 deletions, p = 0.0001, permutation test). The circle highlights the same region in both panels. See text and methods for further details.

Which mechanism can precisely explain the reduced burden of deletions in samples with disrupted common repeat? We propose two non-mutually exclusive scenarios: the “no dissociation scenario” where disruption of the G-quadruplex (GQ) no longer pauses POLG and no longer leads to dissociation, and the “no realignment scenario” where reduced similarity between the two arms of the common repeat decreases the chances of realignment (Figures 1C-1E). To test the “no dissociation” scenario, we analyzed the properties of the GQ that overlaps the first arm of the direct repeat. This GQ sequence is 21 bp long with 3 G-tetrads (positions 8468-8488) and has a pqsfinder (Hon et al., 2017) stability score of 29. Using the pqsfinder tool, we evaluated the impact of two sequence variants, m.8473T>C and m.8472C>T, on the GQ structure. The D4a-specific m.8473T>C variant shortens the GQ to 18 bp but increases its stability to a score of 50. The N1b-specific m.8472C>T variant also reduces the GQ to 14 bp with 2 G-tetrads, decreasing its stability to a score of 10. If GQ length is critical for replication pausing—where longer GQs can pause replication while shorter ones cannot—the shortened versions prompted by both variants suggest that the “no dissociation scenario” remains plausible. The “no realignment scenario” also remains viable as each variant introduces a mismatch, reducing similarity between the arms of the common repeat (see also (Guo et al., 2010)). More research is needed to pinpoint the primary factor that reduces the formation of somatic deletions.

Regardless of the mechanisms that halt deletion formation (whether through “no dissociation,” “no realignment,” or both scenarios), the reduced burden of somatic deletions throughout the body— especially in post-mitotic cells such as neurons and skeletal muscle—in carriers of the disrupted common repeat may contribute to a reduction in age-related mitochondrial diseases and an overall increase in human healthspan. To explore this, we focused on synonymous variants that disrupt the common repeat (D4a-specific m.8473T>C and D5a-specific m.8479A>G), as nonsynonymous variants (like the N1b- specific m.8472C>T) might incur additional fitness costs. Indeed, both synonymous variants, m.8473T>C and m.8479A>G, found in 1% and less than 0.1% of genomes respectively, are associated with longevity [(Bilal et al., 2008), (Alexe et al., 2007), (Tanaka et al. 1998)]. Currently, there is no evidence linking the nonsynonymous N1b-specific variant m.8472C>T with increased lifespan. This association may emerge as more data become available, or the potential benefits from reduced deletions in N1b due to the degraded repeat could be offset by potentially deleterious amino acid substitutions.

Integrating both orthogonal lines of evidence discussed in this chapter—the decreased burden of somatic mtDNA deletions in samples with disrupted common repeats and the association of synonymous SNPs that disrupt the common repeat with increased longevity—we propose that the disrupted common repeat can indeed contribute to healthier aging in carriers by reducing the load of somatic deletions (Figure 1A).

### 4. Disrupted common repeat: merely impacting healthspan or also shaping current and future human evolution?

The disruption of the common repeat introduces an intriguing case where the affected human phenotype (longevity) may not necessarily correlate with evolutionary benefits (fitness). In the exploration below, we delineate three distinct scenarios to understand how this genetic variation could impact evolutionary outcomes: (i) evolutionary neutrality, where the disruption does not affect fitness and thus remains selectively neutral, randomly segregating within the human population; (ii) indirect evolutionary benefits, where increased longevity could indirectly benefit the fitness of subsequent generations, for example, through the grandmother effect; and (iii) direct evolutionary advantages during reproductive years, potentially enhancing fitness through improved health or fertility. We note that the second and third scenarios can coexist and interact; however, each scenario provides a unique lens for examining the impact of disrupted common repeats on human evolutionary trajectories.

This first scenario, “evolutionary neutrality,” suggests that the disruption of the common repeat extends lifespan without impacting fitness, meaning it does not enhance reproductive success or survival. If this is the case, the disruption remains evolutionarily neutral and persists randomly in a population without active selection. However, considering the strong trend towards increased longevity and an extended reproductive age in humans [(Finch, 2010)], this potentially initially neutral mutation, which originated tens of thousands of years ago, might now be accruing indirect (see scenario 2) or direct (see scenario 3) evolutionary benefits. Indeed, the D4a haplogroup and its marker mutation m.8473T>C, which disrupts the common repeat, emerged shortly after the appearance of the D4 haplogroup (28.8-37.1 kya; [(Sukernik et al., 2012)]). At that time, life expectancy was 3-4 times less than the current estimates for the Japanese population, which stands at 81.5 years for males and 86.9 for females [(Organization, 2023)]. This suggests a fascinating period where the selective coefficients of mutations that extend human healthspan are potentially shifting from being effectively neutral at their time of origin to being clearly beneficial now. Additionally, the observed negative correlation between the number of direct repeats in mtDNA and longevity in mammals [(Khaidakov et al. 2006)] further supports the potential evolutionary advantage—both current and prospective—of losing the common direct repeat in humans, a species evolving towards a longer lifespan.

The second scenario, “Indirect Fitness Benefits,” explores indirect fitness benefits, specifically through the grandmother effect, where increased longevity of older generations enhances the survival and fitness of younger family members [(Chapman et al. 2019), (Engelhardt et al. 2019)]. This scenario posits that while the mutation itself—disruption of the common repeat—does not directly enhance the survival rate or reproductive success of the carriers, it does facilitate increased longevity that benefits subsequent generations. Extended lifespans provided by disrupted common repeats might allow grandparents to contribute more significantly to the upbringing and resource provision for grandchildren, potentially increasing the offspring’s chances of survival and reproductive success. The grandmother effect can be particularly pronounced in the case of mtDNA, which is exclusively maternally inherited, aligning evolutionary pressures more closely with female (including grandmother) traits [(Edmands, 2024)]. Altogether, this scenario underscores the existing and growing significance of elder caregiving within contemporary human social structures.

The third scenario, “Direct Fitness Benefits,” suggests that the disrupted common repeat may directly enhance carriers’ survival or reproductive success. The somatic mtDNA deletion load during reproductive age is generally moderate and falls within low-to-moderate heteroplasmy levels, which are expected to be below medically significant thresholds. Thus, this scenario is grounded on the assumption that variations within these low-to-moderate values could potentially impact fitness. For example, 10% instead of 20% of somatic deletion load in postmitotic tissues of carriers versus controls during reproductive age might help maintain these tissues in healthier conditions, reducing sarcopenia and neurodegeneration; similarly, 5% instead of 10% of deletion load in oocytes of carriers versus controls could increase fertilization rates. Although these differences may appear minor, single-cell analyses demonstrate significant heterogeneity within each tissue, indicating that some rare cells could accumulate very high mtDNA deletion load during reproductive years. For instance, analyses of substantia nigra reveal that among neurons typically free of deletions in young samples, a few rare cells are heavily contaminated by deletions (around 60% of heteroplasmy) even in individuals as young as 38 years [(Kraytsberg et al., 2006)]. This suggests a mechanism where a low somatic load at the tissue level corresponds to a high load in some individual cells, which may pass away young, leading to low, but higher than zero levels of sarcopenia, neurodegeneration or other phenotypes, caused by mtDNA deletions. Overall, this scenario suggests that despite low average mtDNA deletion loads during reproductive years, significant heterogeneity at the cellular level may result in minor yet evolutionarily significant variation in phenotypes that could confer slight fitness advantages to carriers. One of such potential weak beneficial effects is considered in more details in chapter 5.

### 5. Mitochondrial genomes of D4a show a decreased rate of molecular evolution

Slight fitness advantages to carriers (hereafter carriers of the common repeat, disrupted by synonymous variants), due to decreased mtDNA deletion load during reproductive age (see scenario 3 in chapter 4), can be challenging to validate experimentally, though such studies are critically needed. However, evolution might detect these subtle effects, for instance, through variations in the mtDNA mutation rate in the germline. We hypothesize that reduced levels of mtDNA deletions in carriers’ oocytes—the sole tissue transmitting mtDNA—might lower the overall germline mtDNA substitutional mutation rate (i.e., the rate of origin of nucleotide substitutions) by reducing background levels of reactive oxygen species (ROS) and/or other damaging factors. Consequently, we suggest that carriers exhibit a lower germline substitutional mutation rate in their mtDNA compared to non-carriers.

The decreased mutation rate, assuming all else is equal, should lead to a reduced rate of nucleotide substitution fixation, which in turn can be observed through shorter branch lengths on the human mitochondrial phylogenetic tree. To evaluate the potential effect of the disrupted repeat on the germline rate of nucleotide substitutions, we utilized our global human tree, reconstructed from more than 43000 complete mitochondrial genomes (see methods). We compared the D4a-specific branch length with its sister clade, sharing the same root. For the branch length estimation, we used several data exclusion thresholds (data having more variability) and a set of substitution matrices ranging from simple to more complex (see methods) because mtDNA is characterized by a very asymmetrical mutation rate with different types of substitutions, sensitive to various life-history traits such as longevity [(Mikhailova et al., 2022)] or basal metabolic rate [(Iliushchenko et al. 2023), (Mikhailova et al. 2023)]. Finally, we demonstrated that D4a, under all models of substitutions and various sequence variability datasets, consistently shows a decreased branch length compared to their background haplogroup (Figure 1F, Supplementary Figure 1, Supplementary Table 2), completely confirming our initial expectations.

The extension of our D4A analyses to other haplogroups with identical mutation m.8473T>C (see Supplementary Table 1) did not reveal any robust trends towards decreased or increased mutation rates in carriers versus non-carriers (Supplementary Table 2). This may be due to either the small sample size of other haplogroups or to a genuine absence of changes in mutation rates and longevity in these groups, reflecting the complex nature of such traits. Drawing an analogy with Liebig’s barrel [(“Liebig’s Law of the Minimum - Wikipedia,” n.d.)], a principle stating that growth is not dictated by total resources available but by the scarcest resource (limiting factor), we can frame our results as follows: a disrupted common repeat can lead to decreased mutation rate and increased longevity if other factors are not limiting this effect. For instance, a disrupted repeat in a haplogroup associated with defective nuclear alleles of POLG or a defective BER pathway would not result in a decreased mtDNA mutation rate and increased longevity. From this perspective, D4a can represent an interesting and important case, where the effect of the disrupted common repeat leads to increased longevity, unimpeded by other nuclear and environmental limiting factors.

## Concluding comments

In this study, we have proposed the causative role of mitochondrial direct nucleotide repeats, particularly the ’common repeat,’ in influencing human healthspan and longevity (Figures 1A). Our analysis has demonstrated that disrupted variants of the common repeat, especially those resulting from synonymous substitutions (Figures 1C-1E), show a promising link to increased longevity due to the decreased burden of somatic deletions which, in parallel with aging, can help mitigate numerous age-related diseases.

Future studies should aim to delve deeper into the anti-aging potential of disrupted common and other repeats, thereby clarifying their therapeutic potential. Examining the distribution and effects of these repeats in diverse human populations, including those in the East (such as haplogroup D4a; Figure 1F), will enrich our understanding of their evolutionary and health-related significance, potentially paving the way for more advanced mitochondrial medicine.

Notably, the likelihood of identifying relevant alleles in aging-related GWAS is low for several reasons. First, mtDNA is often overlooked in these studies [(Melzer et al., 2020)]. Second, even when mtDNA is analyzed, East Asian-specific haplogroups are typically underrepresented in most biobanks [(Yonova-Doing et al., 2021)]. Third, various mtDNA SNPs can reduce the probability of deletion formation, such as mismatches in direct repeats, flanked inverted repeats, or G-quadruplexes (see Figures 1C-1E). This underscores the need to establish an integrated mtDNA metric that approximates mtDNA fragility, akin to a polygenic risk score, to predict the likelihood and frequency of somatic deletions for each mitochondrial genome. When clinically proven, D4a-like alleles could highlight the potential for targeting these mitochondrial mutations either through genome editing or by using mtDNA with disrupted repeats as donor material in mitochondrial transfer or transplantation techniques, as detailed in previous research [(Liu et al. 2022)]. Autologous - derived from the same individual - tissues are currently the primary source for mitochondria isolation in mitochondrial transplantation [(McCully et al. 2023)]. However, using cell cultures as a source provides a readily available supply of mitochondria for immediate use, which could optimize mtDNA for transplantation. This prompts the question of whether D4a-like haplogroups might be the ideal choice for mtDNA in such cell cultures. D4a-like haplogroups with disrupted common repeat might be also considered also as an universal donor of healthy mitochondria in mitochondrial DNA replacement techniques [(Sendra et al. 2021)].

Finally, the described synonymous mutations that disrupt the common repeat are thought to be subject to ongoing weak positive selection in human populations, with the strength of this selection likely to increase in relation to factors such as the advanced age of reproduction. Therefore, it would be interesting to monitor the dynamics of these mutations in the future.

## Methods

### 1. Alignment Preparations

Our study aimed to contrast evolutionary rates of mtDNA genomes encapsulating a major variant of disrupted common repeat with those harbouring both arms of common repeat intact. The analysis hinged on a comprehensive sample of human mtDNA genome sequences sourced from various repositories. To mitigate bias introduced by selection influences, we ensured that the sequences in our sample were devoid of intense selection pressure. This entailed excluding sequences with sites containing deleterious mutations or under significant positive selection, typically rapidly selected during recent mitochondrial evolution, or those with limited variation.

We leveraged the mtDNA genome sequences classification from the HmtDB database (obtained version from October 2018, http://www.hmtdb.uniba.it/) [(Clima et al., 2017)], which segregates mtDNA sequences into two cohorts: those including deleterious mutations and those devoid of such mutations. The multiple sequence alignment with MAFFT (v. 7.4) [(Katoh and Standley 2013)], comprising 43,437 mtDNA genome sequences sans deleterious mutations from HmtDB, served as the foundation for our analysis.

We extracted variable sites from these multiple sequence alignments to ascertain that we analysed neutral or nearly neutral mtDNA sites for the evolutionary rates comparison. This entailed excluding ambiguous characters other than A, T, G or C. We adopted three site variation thresholds: >0·5% (791 sites), >0·1% (1941 sites), and >0·05% (2778 sites) of variation. These thresholds approximately reflect the deviation spectra from the nearly neutral site expectation.

### 2. Phylogenetic Reconstruction

Simultaneously, we predicted a haplogroup for each mtDNA using the HaploGrep v. 2.1.20 software [(Weissensteiner et al., 2016)]. Furthermore, we reconstructed a general Human phylogenetic mtDNA tree using the entire alignment of complete mtDNA genomes with the IQ-TREE v. 1.6.1 software [(Nguyen et al. 2015)]. This employed the general time reversible (GTR) model of base substitutions, along with the FreeRate site variation model [(Soubrier et al., 2012)] allowing for a proportion of invariable sites (option: -m GTR+F+I+R6).

The reconstructed general phylogenetic tree, with marked haplogroups for each sequence, enabled us to clearly demarcate five subsets for the comparative study of evolutionary rates of mtDNA genomes containing disrupted and intact common repeat sequences. These subsets comprised background sequences having normal common repeat arms (control) and foreground sequences manifesting a disrupted common repeat due to the m.8473T>C mutation (case). Using these subalignments, we generated unrooted phylogenetic trees using the IQ-TREE software. Once these unrooted trees were constructed, we utilised the Archaeopteryx software (v. 0.9928 beta) [(Han and Zmasek 2009)] to extract the tree topologies. Tree topology refers to the arrangement of the branches in a phylogenetic tree, which represents the relationships between our five subgroups (Supplementary Figure 2). By extracting the topologies of our trees, we were able to identify the root sequences for each of our subsets. These root sequences represent the common ancestor of the sequences in the subset. Both background and foreground sequences shared a recent common ancestor and were nearly equal in number. With our unrooted trees and identified root sequences, we were then equipped to delve into the analysis of the branch lengths.

### 3. Branch Length Analysis of Evolutionary Rates

Aiming to discern potential disparities in evolutionary rates between our case and control groups, we analysed the branch lengths of the phylogenetic trees generated in the previous phase. A phylogenetic tree’s branch length is a measure of genetic change, and comparing these lengths can provide insights into the evolutionary rates of different groups. However, accomplishing a fair comparison necessitates certain adjustments to account for inherent variances in the data.

The models we selected for our analysis, namely JC, F81, K80, and HKY, are all well-established models of DNA sequence evolution. To clarify, these models describe the probabilities of different types of mutations (such as transitions and transversions) occurring over time. The JC model assumes equal base substitution rates and equal base frequencies, the F81 model implies equal rates but unequal base frequencies, the K80 model involves unequal transition/transversion rates and equal base frequencies, and the HKY model allows for unequal transition/transversion rates and unequal base frequencies.

To root those trees and quantify branch lengths from the root, we used Newick Utilities v. 1.6 [(Junier and Zdobnov 2010)]. Therefore, to ensure an unbiased comparison, we subtracted a value, X/L, from the branch lengths representing the foreground (case) sequences. Here, X is either unity (in the case of JC and F81 models) or the transition/transversion rate (in the case of K80 and HKY models), and L is the number of sites in alignment. This adjustment accounts for the variations in mutation rates and base frequencies that these models inherently consider.

The statistical robustness of our analysis was then tested using a random-half-jackknife procedure (custom Perl script). Jackknifing is a resampling technique used in statistics to estimate the variability of a sample statistic. In our context, the random-half-jackknife procedure involved randomly selecting half of the branches from both the background (control) and foreground (case) groups and comparing their lengths. This process was repeated 100 times to ensure the reliability of our results.

To ascertain the statistical significance of any observed differences in evolutionary rates between the case and control groups, we employed the Wilcoxon signed-rank test (R version 3.4.1; https://www.R-project.org/). This is a non-parametric statistical test that compares two related samples or repeated measurements on a single sample to assess whether their population mean ranks differ. We present results as direction of qualitative shift of U6a, U2e, H1c, R/P and D4 clades with jackknifed mean and standard deviation for p-value quantified by Wilcoxon test for each of F81, HKY, JC and K80 evolutionary models at three variants of site variation stringency selection.

Finally, to visualise the compared background and foreground branch lengths, we employed the vioplot R package v. 0.3.0 [(Adler et al. 2024)], which is particularly useful for comparing distributions across different groups, thus making it apt for our analysis. Through this diligent and comprehensive approach, we ensured a robust and insightful analysis of the evolutionary rates of mtDNA genomes containing disrupted and intact common repeat sequences.

### 4. Global and Local Deletion Patterns Between Samples with Perfect and Disrupted Common Repeats

We analyzed the deletion breakpoint dataset generated using the single-molecule approach performed in [Guo et al., 2010]. This dataset comprises 298 deletions distributed across four samples: 79 deletions in an 82-year-old male with a perfect common repeat, 59 deletions in an 86-year-old male with a perfect common repeat, 85 deletions in an 86-year-old female with a disrupted common repeat, and 75 deletions in a 91-year-old male with a disrupted common repeat. To visualize global and local deletion patterns, all deletions from samples with perfect common repeats were combined and represented in grey in Figure 3, while deletions from samples with disrupted common repeats were combined and shown in red. For the local analysis, we focused on a region centered around the common repeat (13,453 and 8,476 bp) with a radius of 200 bp. In this region, 17 deletions were observed in samples with a perfect common repeat, compared to only 2 deletions in samples with a disrupted common repeat. A permutation analysis was conducted by randomly reassigning the labels (grey or red) of the deletions 10,000 times. The proportion of permutations yielding 2 or fewer red deletions in the circle was 1, resulting in a p-value of 0.0001.

**Figure 3.**
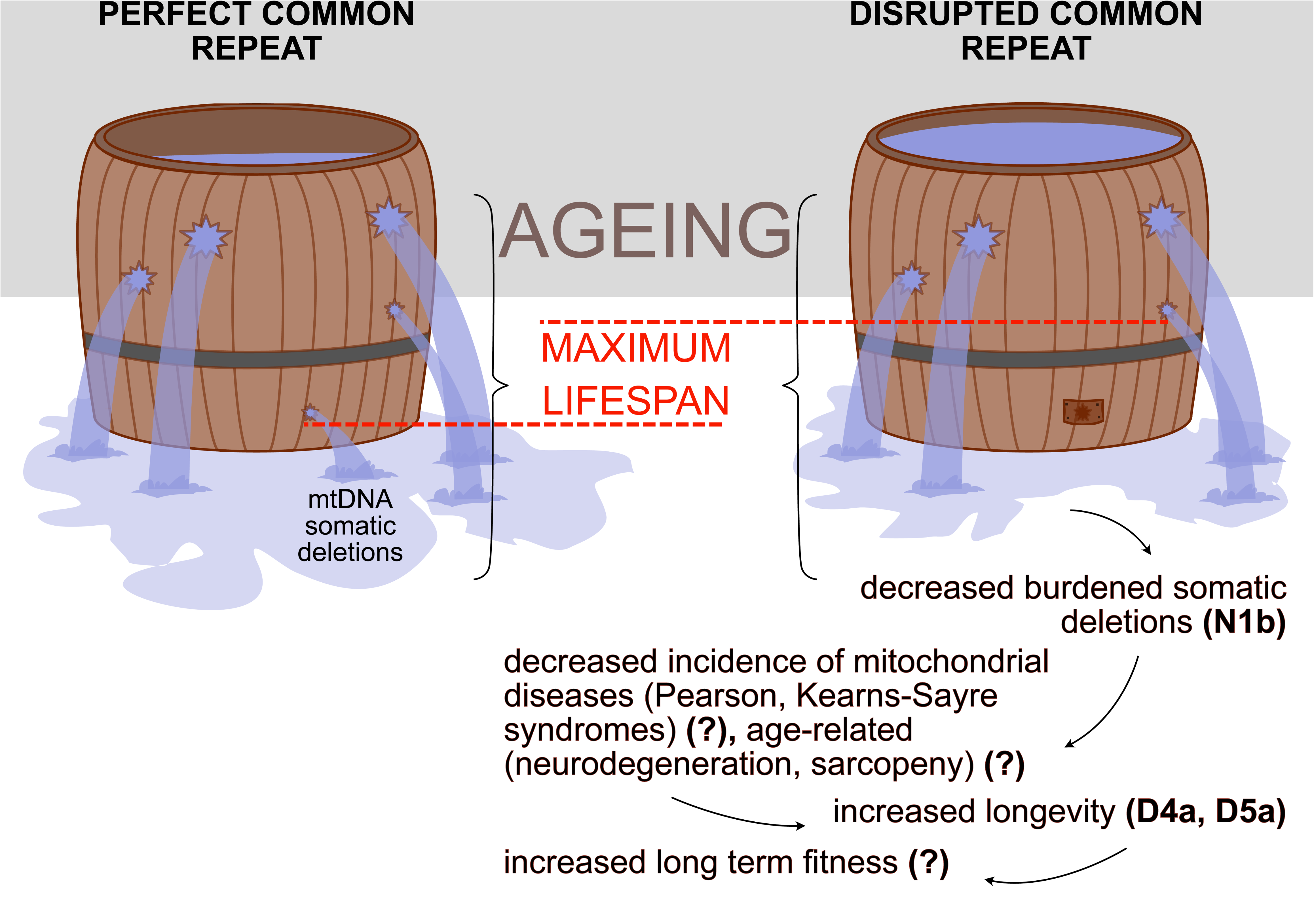
Visual Summary of the working hypothesis: Aging is illustrated as water leaking from Liebig’s barrels, representing the complexity of the aging process. The multiple holes reflect various factors contributing to aging, with mtDNA-related leaks positioned near the bottom due to the lower effective population size of mtDNA compared to nuclear DNA, making it more prone to accumulating all types of slightly deleterious variants including DILL. In the right barrel, a disrupted repeat (fixed hole) slows the water leakage, symbolizing slower aging and a potentially higher maximum lifespan (indicated by a higher water level). Additional phenotypes, some demonstrated and others requiring further investigation (‘?’), are outlined below.

## Availability of data and materials

All data sets and scripts used in the manuscript are available on GitHub https://github.com/mitoclub/JapaneseHypothesis

## Contributions

KP designed the study; KG reconstructed and KG, EOT analysed the phylogenetic tree; VS, EOT, AAM and KP performed main statistical analyses; VS performed an evolutionary analysis of repeats; and all authors wrote the manuscript.

## Ethics declarations

All ethical principles are observed because only anonymised data was used.

## Ethics approval and consent to participate

Not applicable.

## Consent for publication

All co-authors have given their consent to the publication.

## Supporting information

Supplementary Figure 1. The phylogenetic tree of D4a haplogroup (red) with neighbour branches (black). The shortened D4a-specific branches can be exp

Supplementary Figure 2. The phylogenetic tree of haplogroups with m.8473T> C mutation

Supplementary Table 1. The list of variants, disrupting the proximal arm of the common repeat and corresponding haplogroups with more than 20 cases in

Supplementary Table 2. The case-control comparison of the evolutionary rates of mtDNA genomes containing disrupted common repeat (case) with those hav

