## Supplementary figures and images for "Mitochondrial Deletions and Healthspan: Lessons from the Common Repeat"

### Supplementary Figure 1. The phylogenetic tree of D4a haplogroup (red) with neighbour branches (black). The shortened D4a-specific branches can be exp

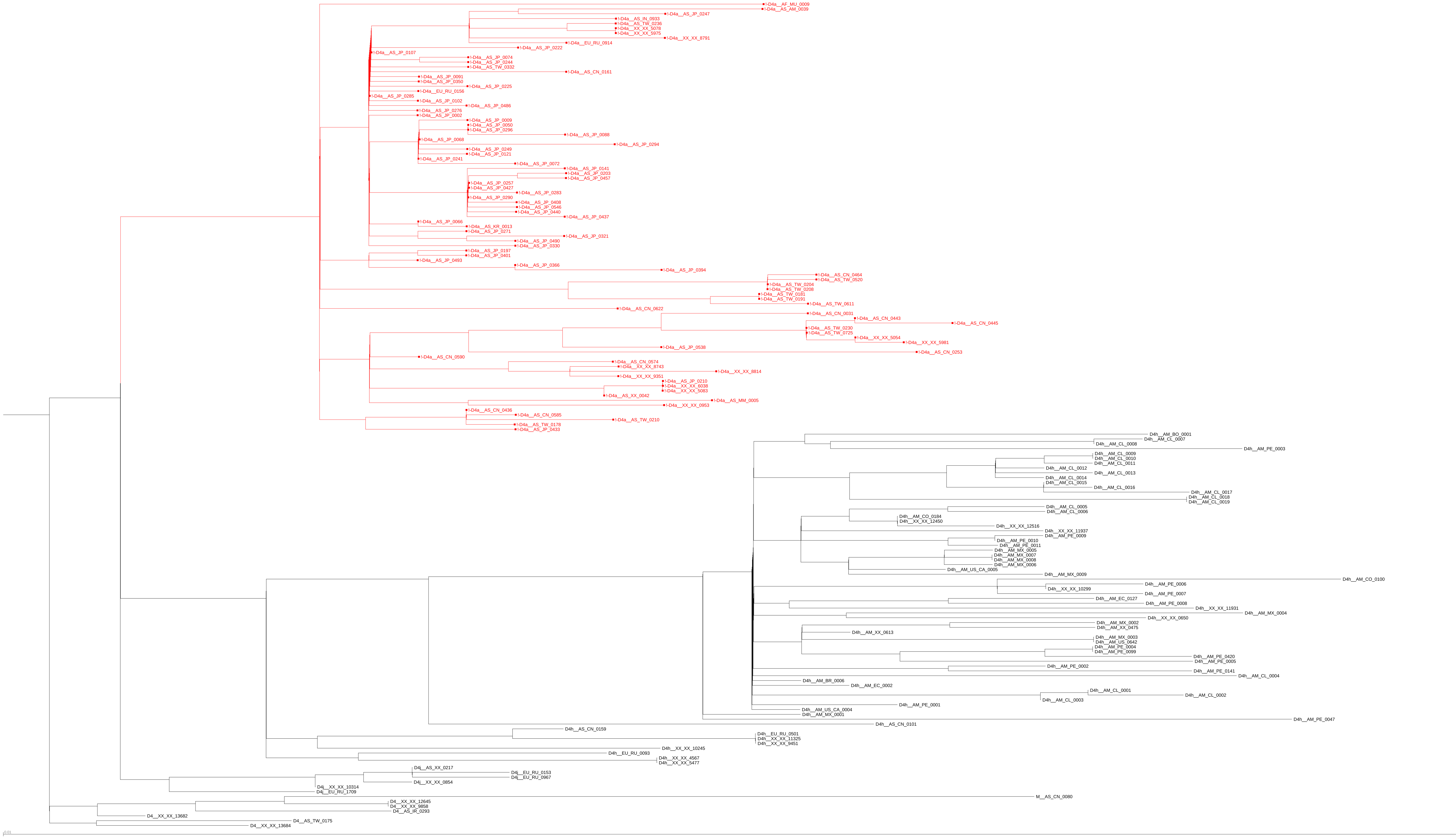

### Supplementary Figure 2. The phylogenetic tree of haplogroups with m.8473T> C mutation

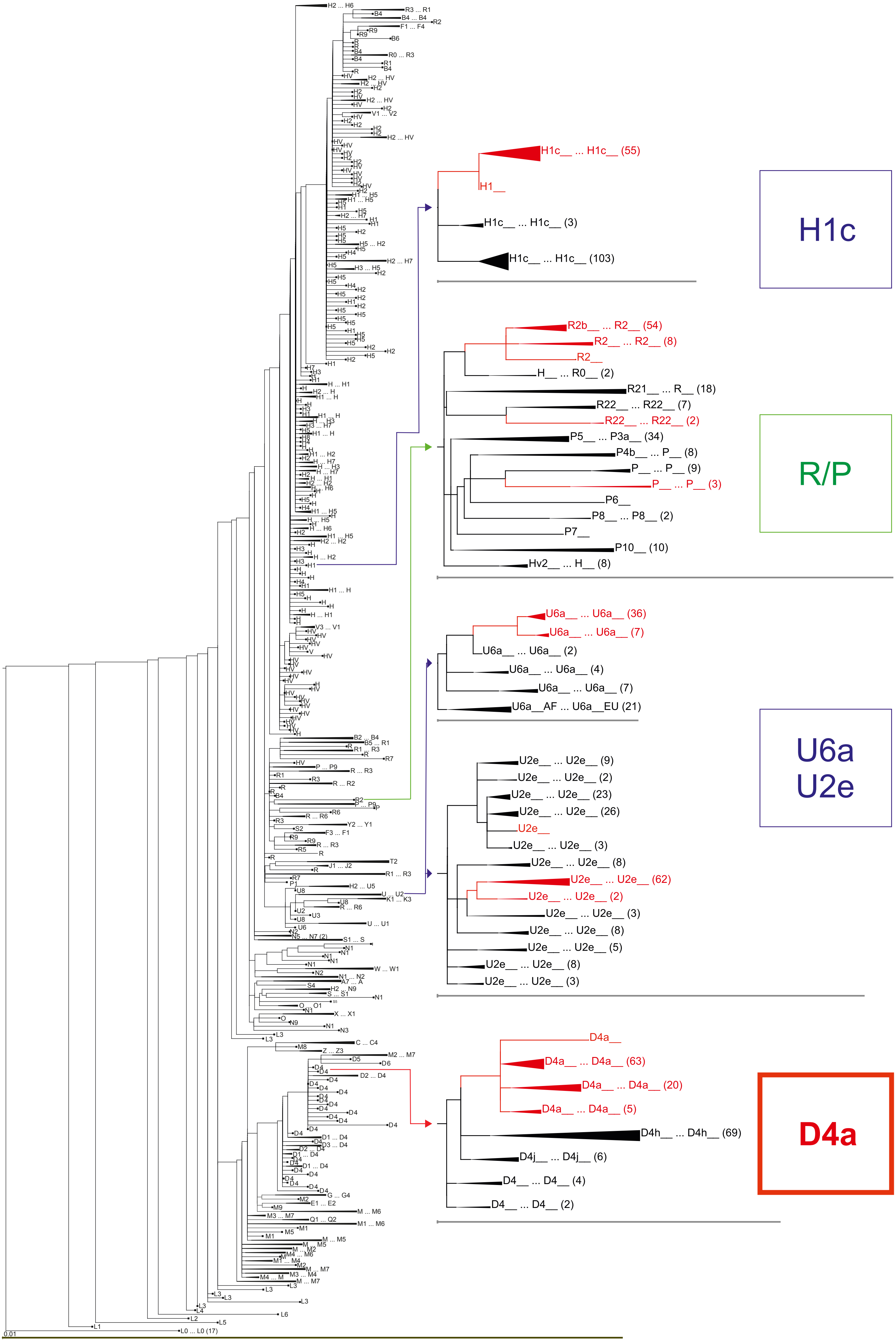
