## Supplementary Table 1. The list of variants, disrupting the proximal arm of the common repeat and corresponding haplogroups with more than 20 cases in for "Mitochondrial Deletions and Healthspan: Lessons from the Common Repeat"

**Supplementary Table 1.** The list of variants, disrupting the proximal arm of the common repeat and corresponding haplogroups with more than 20 cases in HmtDB database (43437 mtDNAs).

| Sequence of the proximal arm of the direct repeat (8470 - 8482 bp) | Haplogroups | Number of individuals |
|---|---|---|
| **the perfect direct repeat: ACCTCCCTCACCA** | **majority of haplogroups, ancestral state and RefSeq** | **42641** |
| m.8473T>C; ACCcCCCTCACCA; SYN | D4a (89), R2 (68), U2e (65), H1c (56), U6a (42), and sporadic cases in other haplogroups (79) | 399 |
| m.8472C>T; ACtTCCCTCACCA; Pro>Leu | N1b (106), and sporadic cases in other haplogroups (38) | 144 |
| m.8479A>G; ACCTCCCTCgCCA; SYN | D5a (57), and sporadic cases in other haplogroups (3) | 60 |
| m.8470A>G; gCCTCCCTCACCA; SYN | Dispersed sporadic cases in various haplogroups (32) | 32 |
| m.8478C>T; ACCTCCCTtACCA; Ser>Leu | L4b (18), and sporadic cases in other haplogroups (12) | 30 |
| m.8477T>C; ACCTCCCcCACCA; Ser>Pro | Dispersed sporadic cases in various haplogroups (26) | 26 |
