## Supplementary Table 2. The case-control comparison of the evolutionary rates of mtDNA genomes containing disrupted common repeat (case) with those hav for "Mitochondrial Deletions and Healthspan: Lessons from the Common Repeat"

**Supplementary Table 2.** The case-control comparison of the evolutionary rates of mtDNA genomes containing disrupted common repeat (case) with those having both arms of common repeat intact (control).

| **Substitution model** | **Sites variation, %** | **Stat. property** | **Clade composition** | | | | |
|---|---|---|---|---|---|---|---|
| | | | **U6a** | **U2e** | **H1c** | **R/P** | **D4** |
| F81 | 0.05 | shift | greater | greater | greater | less | **less** |
| | | average p | 2.17E-05 | 9.73E-10 | 0.00176 | 0.062878 | **7.75E-07** |
| | | std.dev. of p | 5.16E-05 | 1.48E-09 | 0.004146 | 0.091994 | **3.22E-06** |
| | 0.1 | shift | greater | greater | greater | greater | **less** |
| | | average p | 0.00135 | 3.41E-12 | 0.04709 | 0.478095 | **1.23E-06** |
| | | std.dev. of p | 0.001961 | 7.19E-12 | 0.06996 | 0.207913 | **4.38E-06** |
| | 0.5 | shift | less | greater | less | greater | **less** |
| | | average p | 0.198667 | 0.058184 | 0.034564 | 0.281155 | **2.88E-06** |
| | | std.dev. of p | 0.171724 | 0.060021 | 0.033213 | 0.142158 | **6.9E-06** |
| HKY | 0.05 | shift | greater | greater | greater | less | **less** |
| | | average p | 4.27E-05 | 8.09E-12 | 0.04157 | 0.06907 | **2.07E-06** |
| | | std.dev. of p | 9.7E-05 | 1.19E-11 | 0.057705 | 0.091293 | **5.74E-06** |
| | 0.1 | shift | greater | greater | greater | greater | **less** |
| | | average p | 0.005236 | 5.22E-12 | 0.049723 | 0.293431 | **7.91E-07** |
| | | std.dev. of p | 0.013786 | 6.58E-12 | 0.05822 | 0.172256 | **2.48E-06** |
| | 0.5 | shift | less | less | less | greater | **less** |
| | | average p | 0.273149 | 0.284463 | 0.001448 | 0.418316 | **5.53E-07** |
| | | std.dev. of p | 0.209097 | 0.141528 | 0.003522 | 0.236519 | **1.02E-06** |
| JC | 0.05 | shift | greater | greater | greater | less | **less** |
| | | average p | 4E-05 | 6.89E-12 | 0.023705 | 0.0664 | **4.62E-07** |
| | | std.dev. of p | 8.81E-05 | 9.34E-12 | 0.05136 | 0.071869 | **1.22E-06** |
| | 0.1 | shift | greater | greater | greater | less | **less** |
| | | average p | 0.000618 | 2.35E-11 | 0.044148 | 0.5305 | **2.71E-07** |
| | | std.dev. of p | 0.001676 | 4.55E-11 | 0.062387 | 0.214982 | **6.51E-07** |
| | 0.5 | shift | less | greater | less | less | **less** |
| | | average p | 0.039706 | 0.237216 | 0.009861 | 0.461538 | **8.35E-06** |
| | | std.dev. of p | 0.060181 | 0.186655 | 0.014167 | 0.266469 | **1.69E-05** |
| K80 | 0.05 | shift | greater | greater | greater | less | **less** |
| | | average p | 9.73E-06 | 1.66E-11 | 0.031127 | 0.087671 | **1.14E-07** |
| | | std.dev. of p | 7.1E-06 | 5.27E-11 | 0.025872 | 0.146845 | **2.04E-07** |
| | 0.1 | shift | greater | greater | greater | greater | **less** |
| | | average p | 0.000228 | 6.21E-10 | 0.060003 | 0.369783 | **3.65E-07** |
| | | std.dev. of p | 0.000542 | 1.61E-09 | 0.106046 | 0.204312 | **1.19E-06** |
| | 0.5 | shift | less | greater | less | greater | **less** |
| | | average p | 0.017182 | 0.189886 | 0.02839 | 0.554828 | **3.98E-06** |
| | | std.dev. of p | 0.019592 | 0.11964 | 0.043765 | 0.255987 | **9.78E-06** |
